# T cells modulate the microglial response to brain ischemia

**DOI:** 10.1101/2021.09.23.461546

**Authors:** Corinne Benakis, Alba Simats, Sophie Tritschler, Steffanie Heindl, Simon Besson-Girard, Gemma Llovera, Kelsey Pinkham, Anna Kolz, Fabian Theis, Özgün Gökce, Anneli Peters, Arthur Liesz

## Abstract

Neuroinflammation after stroke is characterized by the activation of resident microglia and the invasion of circulating leukocytes into the brain. Although lymphocytes infiltrate the brain in small number, they have been consistently demonstrated to be the most potent leukocyte subpopulation contributing to secondary inflammatory brain injury. However, the exact mechanism how this minimal number of lymphocytes can profoundly affect stroke outcome is still largely elusive. Here, using a mouse model for ischemic stroke, we demonstrated that early activation of microglia in response to stroke is differentially regulated by distinct T cell subpopulations. Acute treatment with engineered T cells overexpressing IL-10 administered into the cisterna magna after stroke induces a switch of microglial gene expression to a profile associated with pro-regenerative functions. These findings substantiate the role of T cells in stroke with large impact on the cerebral inflammatory milieu by polarizing the microglial phenotype. Targeting T cell-microglia interactions can have direct translational relevance for further development of immune-targeted therapies for stroke and other neuroinflammatory conditions.

**Summary:** The crosstalk between brain infiltrating T cells and microglia in response to stroke remains elusive. Benakis et al. report that transcriptional signature of the stroke-associated microglia is reprogrammed by distinct T cell subpopulations. Engineered T cells overexpressing IL-10 administered four hours after stroke reinitiate microglial function inducing a pro-regenerative environment.

## Introduction

Among peripheral leukocytes invading the injured brain, T cells have been consistently identified as the invading leukocyte subpopulation with the largest impact on secondary neurodegeneration and modulation of the ischemic brain damage (Kleinschnitz et al., 2010; Liesz et al., 2011, 2009). T cell subpopulations have the potential to play either a neuroprotective or a deleterious role in post-stroke neuroinflammation. In particular, the pro-inflammatory T_H1_, T_H17_ subsets of T_HELPER_ cells and IL-17-producing γδ T cells have been shown to induce secondary neurotoxicity, leading to infarct expansion with worse functional outcome (Gelderblom et al., 2012; Shichita et al., 2009), whereas regulatory T cells (T_REG_) exert anti-inflammatory and neuroprotective function suppressing an excessive inflammatory reaction to the brain infarct (Liesz et al., 2009). Recruitment of peripheral immune cells is not limited to the brain parenchyma since an accumulation of T cells is observed in the choroid plexus and the meninges after stroke (Benakis et al., 2016; Llovera et al., 2017). Attempt in blocking the recruitment of peripheral effector T cells diminished neuronal damage in different cerebral ischemic models, resulting in improvement of stroke outcome and suggesting a possible therapeutic target (Liesz et al., 2011; Llovera et al., 2015). Considering the relatively low number of only a few thousand lymphocytes invading the brain after stroke compared to more than fifty times higher cell count of innate immune cells (invading and resident) in the post-stroke brain (Gelderblom et al., 2009), it is surprising to observe such a dramatic effect of a small number of T cells on the neuroinflammatory response to stroke.

Therefore, we hypothesized that T cells have a polarizing effect on microglial function. In turn, microglia – as the most abundant immune cell population in the ischemic brain – could amplify the T cells’ impact on the cerebral immune milieu. Indeed, microglia interact with T cells via either cell-to-cell contact, cytokine-mediated communication or antigen presentation leading to activation/polarization of adaptive immune cells entering the brain (Goldmann and Prinz, 2013). T cell–microglia interaction can further influence the neuroinflammatory response in experimental models of multiple sclerosis (Dong and Yong, 2019) and possibly in stroke (Wang et al., 2016). In fact, recent evidence suggest a crosstalk between microglia and T cells as a key determinant of neuronal plasticity during recovery from brain injury (Shi et al., 2021). However, while the influence of microglia/macrophages on T cells has been well-studied, it is still unclear how in reverse the T cells influence microglial function, and whether early interaction of T cells with microglia in the acute response to stroke can have an immediate impact on microglia and further change the course of disease progression.

Using single-cell sequencing and adoptive transfer models of *ex vivo* differentiated T_HELPER_ cell subpopulations, we performed an in-depth analysis of the immunomodulatory effects of T cells on microglial polarization. Better understanding of the T cell–microglia crosstalk holds the potential to use polarized T cells as a therapeutic approach with large impact on the cerebral inflammatory milieu potentiated by resident microglia.

## Results

### Lymphocytes modulate the activation state of microglia in response to stroke

First, we investigated the effect of lymphocytes on microglial morphology and transcriptome in lymphocyte-deficient Rag1^-/-^ mice after experimental stroke (**Fig. 1a**). Microglia were analyzed 5 days after stroke because this acute time point has previously been identified as the time of maximal cerebral lymphocyte infiltration in the same stroke model (Llovera et al., 2017). Using an automated morphological analysis of IBA1 positive cells (Heindl et al., 2018), we identified that microglia in the perilesional (ipsilateral) cortex of Rag1^-/-^ mice displayed extended ramifications and a lower sphericity compared to microglia of wild type (WT) mice, indicating a less activated microglial phenotype in the absence of lymphocytes (**Fig. 1b, c**). In contrast, microglial morphology remained unchanged between Rag1^-/-^ and WT mice in the contralateral (unaffected) hemisphere which does not show recruitment of lymphocyte in considerable amounts, supporting the role of local lymphocyte infiltration for changing microglial morphology. Therefore, we investigated the functional implications of cerebral lymphocyte invasion for microglia by single-cell transcriptomics in WT and Rag1^-/-^ mice. CD45^+^CD11b^+^ myeloid cells were sorted by flow cytometry from naïve mice or 5 days after stroke (pool of 3 mice per condition) and single-cell RNA sequencing was performed using 10x Genomics (**Fig. 1d**). Unsupervised clustering analysis identified 14 distinct clusters across conditions (**Fig. 1e** and Supplementary Fig. 1a). Based on the expression of previously defined markers of homeostatic and reactive microglia per cell cluster (gene expression of *Fcrls, P2ry12, Trem2*; absence of *Lyz2, Ccr2*; (Keren-Shaul et al., 2017; Miron and Priller, 2020; Prinz and Priller, 2014)), 5 clusters were annotated as microglial cells (**Fig. 1e, right**, and Supplementary Fig. 1a). We then further clustered the microglial cells into subpopulations showing either a transcriptomic profile associated with homeostatic microglial function (clusters 0, 1, 4, 6) or a profile of reactive microglia, activated in response to the ischemic tissue injury (clusters 2, 3, 5, 7) (Supplementary Fig. 1b, c). The cell distribution across condition highlighted that stroke induced a reactive transcriptomic profile in the majority of microglia derived from the perilesional cortex, both in WT and Rag1^-/-^ mice (**Fig. 1f**).

**Figure 1.**
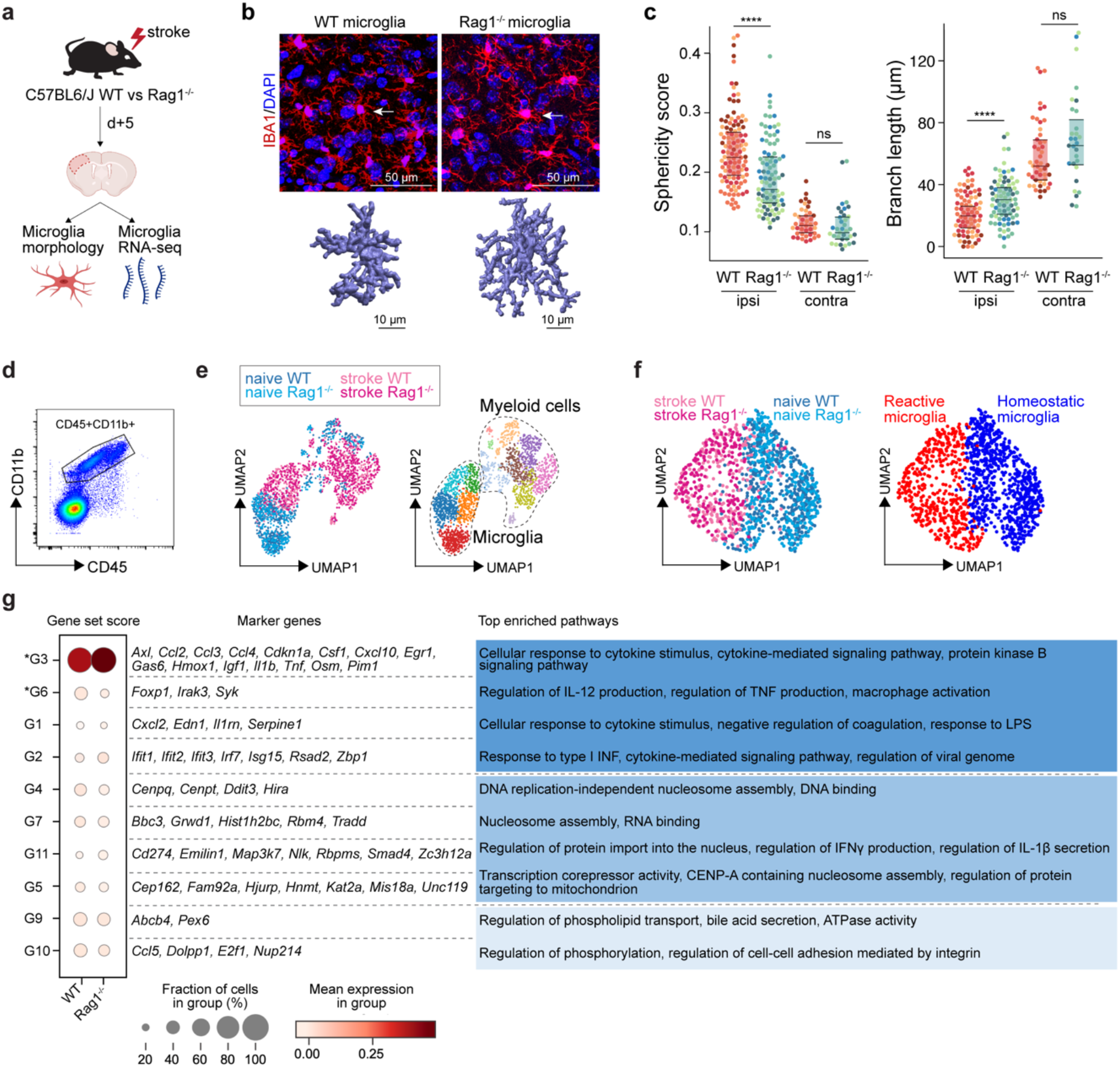
T cells influence microglia morphology and transcriptomic signature. **a** Morphological analysis of microglia and transcriptomic profile of sorted microglia were performed 5 d after stroke in wild-type (WT) and Rag1^-/-^ mice. **b** Top, representative images of IBA1+ microglial cells in the perilesional region (900 μm distal to the infarct border, cortical layer 4). Bottom, threedimensional (3D) reconstruction of microglia in WT and Rag1^-/-^ mice. **c** Morphological analysis of microglia in the peri-infarct area (ipsi) and in the contralateral hemisphere (contra) for two representative features: sphericity and branch length (μm) in WT (orange) and Rag1^-/-^ (blue) mice. Each individual mouse is represented in the plots by one color (4 mice/condition), each dot corresponds to one microglial cell; ns, non significant; ****, P < 0.0001. Wilcoxon rank sum test with continuity correction and Bonferroni post-hoc correction for multiple testing. **d** CD45+CD11b+cells were sorted from the ipsilateral hemisphere 5 d after stroke in WT and Rag1^-/-^ (3 mice/condition) and RNA was isolated for single cell RNA sequencing (10x Genomics). **e** Uniform manifold approximation and projection 2D space (UMAP) plots of 2345 CD45+CD11b+ cells colored by conditions (left) and by 14 distinct transcriptional clusters (right and Supplementary Fig. 1). **f** Clustering of the microglia subset color-coded by conditions (left) and into homeostatic and reactive microglia (right). **g** Selected gene sets of highly correlated and anti-correlated genes based on trajectory inference analysis in stroke condition (Supplementary Fig. 1d–f). Mean gene set activation score in WT and Rag1^-/-^ cells, selected marker genes and top enriched gene ontology pathways associated to each gene set. Gene sets were classified by p-value (the lowest p-value at the top, asterisks (*) indicate significant difference between genotype in stroke condition) and by similar pathways, such as: pathways related to inflammation (dark blue), pathways related to DNA/RNA regulation (blue) and lipid pathways (light blue).

The microglial reaction to stroke causes a gradual shift from the homeostatic transcriptomic profile to a reactive state. In order to capture differences in the microglia transcriptome along its transition phase, we performed single cell trajectory inference analysis (Supplementary Fig. 2). Partition-based graph abstraction (PAGA) revealed two distinct paths with high connectivity from the homeostatic (naïve) microglia cluster (root cluster) to the reactive (stroke) microglia cluster (end cluster) (Supplementary Fig. 2a). Interestingly, the number of microglial cells in stroke Rag1^-/-^ increased in the end cluster and was decreased along the trajectory path 2 in comparison to stroke WT (Supplementary Fig. 2b), suggesting that lymphocytes influence the transition of a microglia subpopulation from the homoeostatic to the reactive state. Differential gene expression analysis between the root and end clusters of the trajectory path 2 in WT and Rag1^-/-^ mice (Supplementary Fig. 2c) revealed that genes associated with ribosomal metabolic processes and mitochondrial ribosomal proteins were specifically enriched whereas genes associated with phagocytosis were down-regulated in microglia of Rag1^-/-^ mice (Supplementary Fig. 2d). We then clustered genes into groups of correlating and anti-correlating genes and investigated the activation of these gene sets along the identified trajectory path 2 in stroke condition only (Supplementary Fig. 1d–f). Gene sets which were significantly different between WT and Rag1^-/-^ mice after stroke revealed that the absence of lymphocytes significantly reduces microglial genes associated with macrophage activation state (G6: *Foxp1, Syk*), whereas genes associated with cytokine/chemokine stimulus were enriched (G3: *Il1b, Tnf, Csf1, Ccl2*) in Rag1^-/-^ in comparison to WT microglia (**Fig 1g**).

Next, we aimed to validate our scRNA-seq findings by an independent transcriptomic platform using Smart-seq2 profiling of sorted pools of 50 microglia which provides cell type specificity and higher mRNA capture than scRNA-seq (Picelli et al., 2014). We confirmed a predominant effect of stroke on the microglial transcriptome (**Fig. 2a, b**), in which lymphocyte-deficiency affected the abundance of specific microglia subsets (**Fig. 2c–e**). In order to identify the pathways specifically regulated in these subsets, we performed a ‘Sample-Gene cluster’ correlation analysis and identified two gene sets that were significantly regulated between WT and Rag1^-/-^ mice after stroke (i.e., G1 and G3; **Fig. 2f, g**). Specifically, we detected a decrease of gene expression associated with glial cell migration and leukocyte differentiation (gene cluster G3), and an increase in neutrophil degranulation and cytokine production (G1) in lymphocyte-deficient mice (**Fig. 2g**), supporting our results from the single cell sequencing obtained with the 10x protocol. Altogether, we demonstrated that lymphocytes modulate the activation status of a subset of stroke-associated microglial cells towards a phenotype associated with increased phagocytosis and immune cell accumulation in the post-ischemic brain.

**Figure 2.**
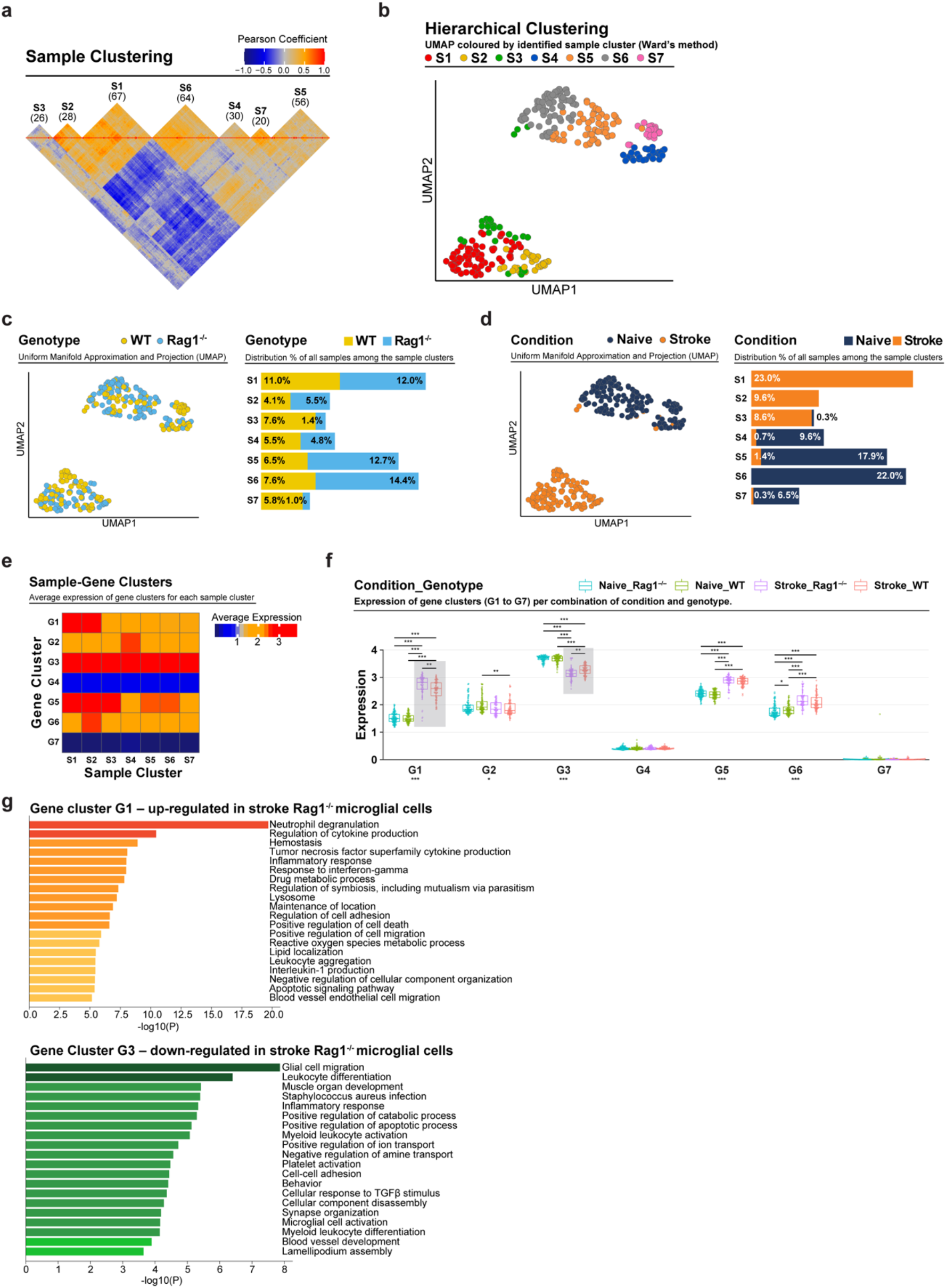
Analysis of microglia isolated from WT and Rag1^-/-^ in naïve and stroke conditions using the Smart-Seq2 platform. **a** Correlation between the 291 pools of microglia. Hierarchical clustering identified 7 clusters, named from S1 to S7. The amount of samples is in between parentheses. The 1000 most variable genes were considered. The color represents the Pearson correlation coefficient between the samples. **b–d** UMAP plots showing the samples colored by clusters (b), genotype (c) and condition (d) and their associated cell distribution of genotypes and conditions among the sample clusters. **e** Average expression of gene clusters (defined in panel g) within each sample cluster. Scale is the average of the log normalized expression. **f** Box plots of the gene clusters in each conditions and genotypes. Grey boxes highlight significant difference between WT and Rag1^-/-^ in stroke condition. *, P < 0.05; **, P < 0.01; ***, P < 0.001. Nonparametric Kruskal Wallis test followed by multiple non-parametric Wilcoxon t-tests and Bonferroni post-hoc correction for multiple testing. **g** Gene clusters and associated pathway enrichment analysis. Hierarchical clustering identified 7 gene clusters, named G1 to G7, only the significantly regulated gene clusters G1 and G3 between WT and Rag1^-/-^ in stroke condition are shown. Each barplot shows the pathway enrichment analysis for the genes included in the gene clusters.

### T_HELPER_ cell subpopulations drive the distinct polarization of microglia

To test whether microglial phenotypes can be specifically skewed by functionally opposing T_HELPER_ cell subpopulations, we differentiated T_H1_ and regulatory T cells (T_REG_) *in vitro* (Supplementary Fig. 3a) and tested whether these T_HELPER_ cells can reprogram the stroke-associated microglia. Differentiated T cells or vehicle were injected into the cisterna magna (CM) of lymphocyte-deficient Rag1^-/-^ mice 24 h after stroke. Microglia cells were sorted from the ipsilesional hemisphere 24 h after polarized T_HELPER_ cell (T_H1_ or T_REG_ cells) or vehicle administration (**Fig. 3a**). The transcriptional profile of microglia induced by T_REG_ cells was more similar to vehicle treated Rag1^-/-^ mice (named control (CT)), than microglial gene expression induced by T_H1_, as shown in the heatmap and volcano plots of the differentially expressed genes (P < 0.05 and |fold change| > 3) with 51 and 20 microglial genes regulated in T_H1_ or T_REG_ conditions compared to control injection, respectively (**Fig. 3b, c)**. Gene ontology analysis of the differentially up-regulated genes revealed T_H1_-dependent pathways associated with antigen presentation, response to cytokines and regulation of type I interferon whereas T_REG_-dependent microglial genes were associated with chemotaxis **(Fig. 3d)**. These results demonstrate the potency of T cell subpopulations to differentially skew the microglial transcriptome towards distinct phenotypes previously associated with different cellular functions. In particular, we found that T_H1_ polarized microglia toward an antigen-immunocompetent phenotype (*Cd74*) and expression of interferon response-related genes (*Irf7*). This profile of microglial response was previously associated with a pronounced immune response during the later stages of neurodegeneration (Mathys et al., 2017). In contrast, T_REG_ cells promoted the expression of chemokines/cytokines in microglia (*Ccl2, Ccl7, Cxcl10*), which can have either pro-regenerative or detrimental effects such as the regulation of leukocyte chemotaxis to the injured brain (Llovera et al., 2017), mechanisms of protective preconditioning (Garcia-Bonilla et al., 2014) or promoting neuronal stem cell recruitement and angiogenesis (Andres et al., 2011; Lee et al., 2012; Liu et al., 2007). In addition, after experimental stroke, the T_H1_-mediated effects on the microglial transcriptomic profile were associated with an increase of *Trem2* expression, a key marker of disease-associated microglia in various brain disorders, in comparison to microglia primed by T_REG_ cells (**Fig. 3e**). These transcriptomic differences in microglia related to the *in vivo* T_H1_ or T_REG_ cell exposure was also reflected by the difference in the morphology of microglia between these conditions. Microglia displayed a reactive state as shown by a more spherical and less branched morphology in T_H1_ cell-injected compared to T_REG_-injected mice (**Fig. 3f**).

**Figure 3.**
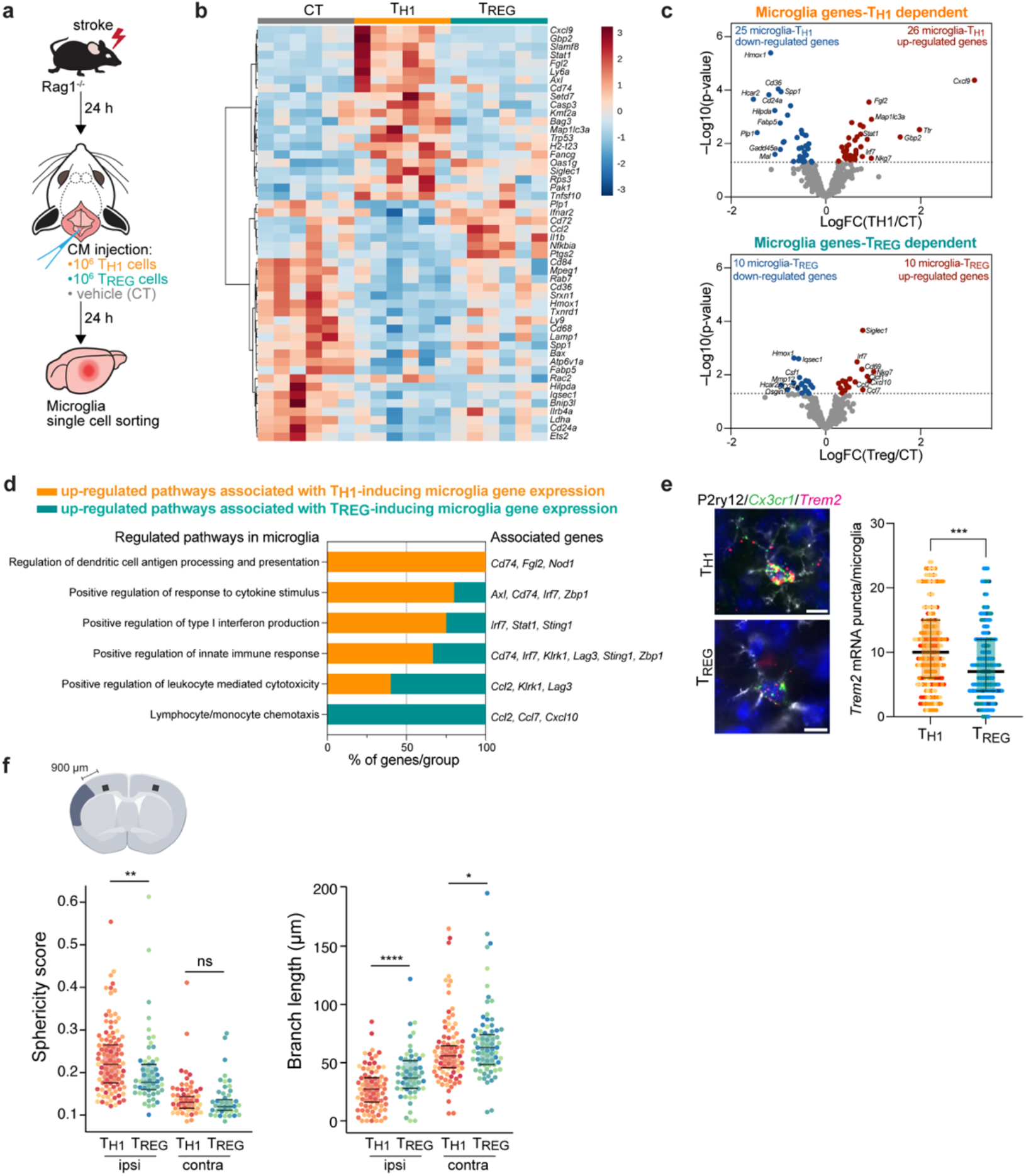
T_H1_ and T_REG_ cells influence microglia gene expression after stroke. **a** Naïve CD4 cells were polarized in vitro to T_H1_ or T_REG_ phenotype (Supplementary Fig. 3a). One million cells (T_H1_ or T_REG_ cells) or vehicle (control, CT) were injected into the cisterna magna (CM) in Rag1^-/-^ mice 24 h after stroke induction. Microglia cells were sorted from the ipsilesional hemisphere and RNA was extracted. Gene expression analysis was performed using the Neuroinflammation Panel profiling kit on the Nanostring platform. In a second set of experiment, 100 μm coronal sections were proceeded for smFISH or microglia morphology. **b** Heatmap representation of microglia gene expression between conditions: control (CT; vehicle administration of PBS), T_H1_ or T_REG_. **c** Up- and down-regulated differentially expressed genes between either isolated microglia from T_H1_- (top) and T_REG_- (bottom) treated Rag1^-/-^ mice relative to control condition (microglia isolated from Rag1^-/-^ mice treated with vehicle, genes are color-coded accordingly to a p-value < 0.05 and |fold change| > 3). **d** Pathway analysis was performed for the up-regulated genes in each condition using the ClueGO package from Cytoscape. **e** Higher amount of *Trem2* mRNA puncta (red) per *Cx3cr1*-positive (green) in P2ry12-labelled microglia (white) in T_H1_-treated mice in comparison the T_REG_-treated mice. DAPI (blue) was used as nuclear dye. Scale bar = 10 μm. **f** Morphological analysis of IBA1+ microglia in the ipsilateral (900 μm distal to the infarct border, cortical layer 4) and contralateral hemisphere, as shown in the representative coronal section. Sphericity score and branch length (μm) of microglia treated with T_H1_ (orange) or T_REG_ cells (green). Each individual mouse is represented in the plots by one color (3 mice/condition), each dot corresponds to one microglial cells; ns, non significant; *, P < 0.05; **, P < 0.01; ****, P < 0.0001. Wilcoxon rank sum test with continuity correction and Bonferroni post-hoc correction for multiple testing.

Interestingly, these morphological changes were not only restricted to the ipsilesional hemiphere but were also observed in the contralateral hemisphere, suggesting possible brain-wide effects of differentiated T_HELPER_ cells injected to the cerebrospinal fluid (CSF) compartment. In accordance, we found that intra-CM injection of eGFP-labelled T_H1_ cells to Rag1^-/-^ mice after stroke were primarily recruited to the ischemic brain parenchyma, but were additionally also localized in border tissues including the meninges, and some CM-injected cells even circulated and could be detected in the spleen. (**Fig. 4a, b** and Supplementary Fig. 3b). Together, these findings support that polarized T cells are recruited to the infarction site and may modify *in situ* the inflammatory micromilieu.

**Figure 4.**
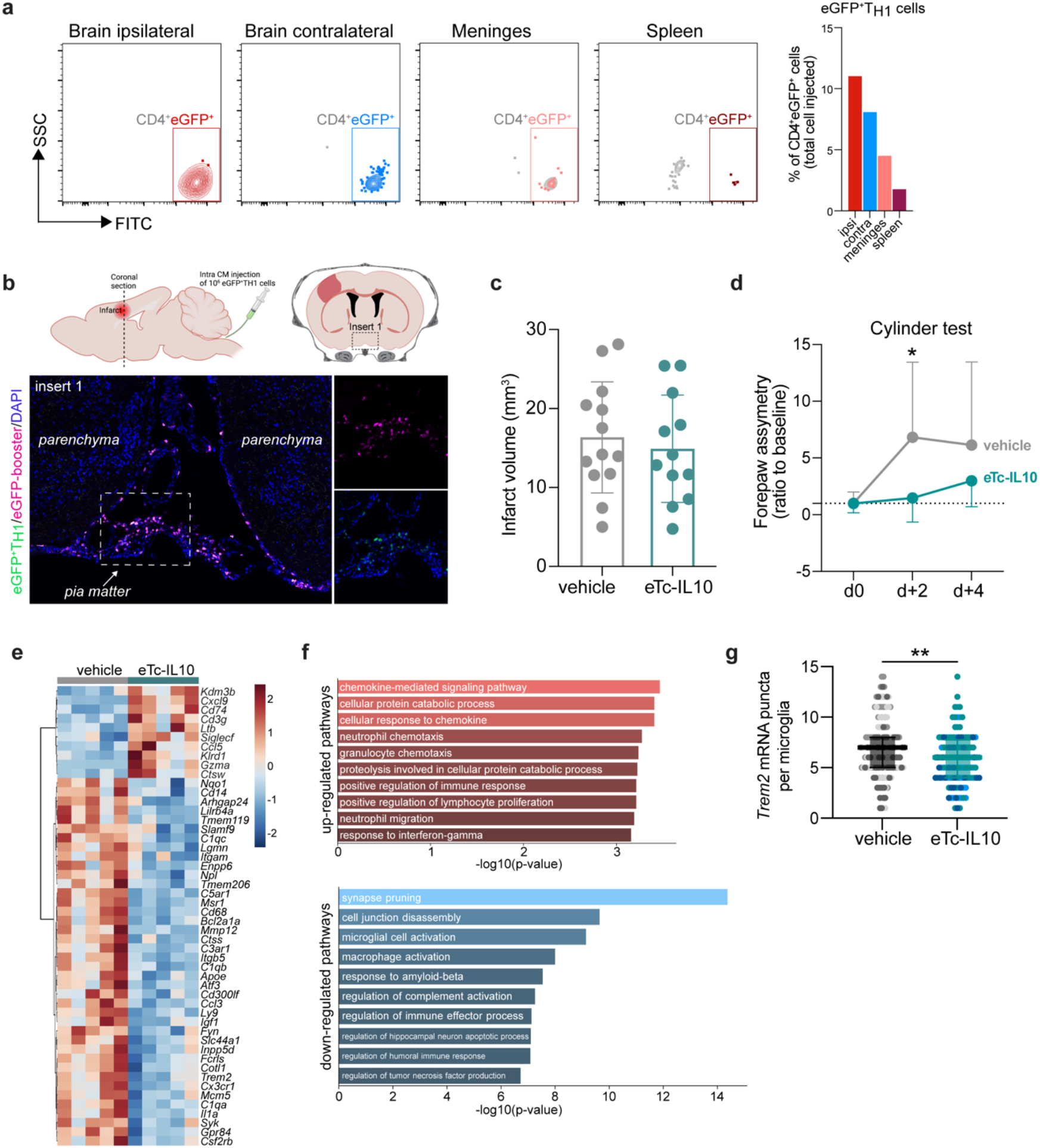
Acute post-stroke treatment with engineered T cells overexpressing IL-10 modulates microglial activation and ameliorates functional deficit. **a, b** Flow cytometry analysis and whole skull-brain coronal sections of 10^6^ eGFP+T_H1_ cells injected into the cisterna magna (CM) of Rag1^-/-^ mice 24 h after stroke. Samples were collected 4 h after CM injection for further analysis: **a** Flow cytometry plots showing CD4+eGFP+ cells isolated from the brain (ipsilateral and contralateral hemispheres), meninges and spleen (the detailed gating strategy is shown in Supplementary Fig. 3b). The graph represents the percentage of eGFP+T_H1_ cells relative to the total number of cells injected in the CM (10^6^ eGFP+T_H1_ cells). **b** Coronal section showing eGFP+T_H1_ cells in the meninges. Insert 1 indicates a representative photomicrograph of eGFP+T_H1_ cells counterstained with an eGFP-booster (magenta), cell nuclei are stained with DAPI (blue). The magnified images of white boxed area show eGFP+T_H1_ cells injected into the CM are located in the meninges. **c** Infarct volumes at 5 d after stroke in WT C57BL/6J mice treated by CM administration of either T cells secreting IL-10 (eTc-IL10, 10^6^ naïve CD4+ cells transfected with a plasmid overexpressing IL-10, Supplementary Fig. 3c, d) or vehicle (aCSF) 4 h after stroke induction. **d** Percentage of assymetry in independent forepaw use (“0%” indicates symmetry) in mice treated with vehicle or eTc-IL10; *, P < 0.05, ANOVA with Šídák’s multiple comparisons test. **e** Heatmap representation of ipsilateral brain gene expression between vehicle and eTc-IL10 treated mice 5 d after stroke. **f** Selected gene ontology annotations for the 50 genes that were up-regulated (top) and down-regulated (bottom) in the whole ipsilateral brain tissue of eTc-IL10 treated mice in comparison to vehicle treated mice. **g** smFISH analysis of brains from eTc-IL10 treated mice showed a reduction of *Trem2* mRNA puncta per *Cx3cr1*-positive microglia in the peri-infarct region in comparison to vehicle treated mice; **, P < 0.01, Mann-Whitney U test.

### Engineered T cells overexpressing IL-10 induce a pro-regenerative microglial phenotype

In order to further explore the therapeutic potential of T cell-secreted cytokines to modulate the local microglial immune milieu, we engineered T cells by viral transfection to overexpress the anti-inflammatory cytokine IL-10 (eTc-IL10; Supplementary Fig. 3c,d). In a therapeutic approach, we injected eTc-IL10 cells into the CM of WT mice 4 h after stroke – a translationally relevant time window considering a similar time window for acute therapy with thrombolytics in stroke patients. We investigated whether eTc-IL10 treatment affected stroke outcome but did not find any difference in infarct volumes between conditions **(Fig. 4c**). This is in accordance with the concept of early ischemic lesion formation in stroke which is not being affected by the delayed immunological mechanisms (Dirnagl et al., 1999). In contrast, mice receiving eTc-IL10 injection in the CM had a substantially improved functional outcome at 48 h after stroke as shown by a reduced forelimb assymetry (**Fig. 4d**). This might reflect the particular impact of inflammatory pathways and specifically cytokine secretion on functional deficits and delayed recovery after stroke in contrast to the early primary lesion development (Filiano et al., 2017; Roth et al., 2020). We then evaluated whether gene expression was altered after stroke upon eTc-IL10 treatment. RNA was isolated from the whole ischemic hemisphere and neuroinflammatory genes were quantified using the Nanostring platform. Interestingly, we found that several genes associated with a disease-associated microglial profile were down-regulated in mice treated with eTc-IL10 such as *CD68, Apoe, Itgax, Trem2, Tyrobp and Cst7* (**Fig. 4e** and Supplementary Fig. 3e). Gene ontology analysis revealed that T cell-derived IL-10 overexpression increased pathways associated with chemokine responses, and the downregulation of several microglial effector functions such as spine pruning, phagocytosis and complement activation **(Fig. 4f)**. This anti-inflammatory effect of eTc-IL10 treatment on microglia was confirmed by a reduction of *Trem2* mRNA in *Cx3cr1+* microglia from eTc-IL10 compared to vehicle-treated mice (**Fig. 4g**). Since we observed a down regulation of genes associated with synapse pruning (*C1qa, C1qc*), microglia activation (*P2ry12, Cx3cr1*) and phagocytosis (*Trem2*) in mice treated with eTc-IL10, we postulate that acute intra-CM administration of eTc-IL10 induces a switch of genes characteristic of homeostatic microglia possibly promoting post-stroke recovery mechanisms.

## Discussion

The cellular constituents of the acute neuroinflammatory response to stroke has been well characterized including microglial activation, leukocyte invasion and the contribution of different lymphocyte subpopulation (Anrather and Iadecola, 2016). However, the reciprocal interactions of these different immune cell populations remain largely underinvestigated in the context of brain injury. A better understanding of the T cell polarizing effect on microglial function has strong translational implication, since T cells may act as a ‘Trojan horse’ with large impact on the cerebral inflammatory milieu potentiated by microglia (Cramer et al., 2018).

Here, we established a mechanistic link between T cells and microglial function and showed the distinct role of T cell subpopulations on switching microglial polarization state in response to stroke. Our results from transcriptomic analysis suggest that the microglia-polarizing effect of different T_HELPER_ cell subpopulations is mainly mediated via their specific cytokine secretion pattern. Microglia that were challenged with T_H1_ cells expressed an upregulation of genes associated with type I interferon signaling (INF) – the key cytokine secreted by the T_H1_ subpopulation. In contrast, T_REG_ cells modulated a gene set associated with chemotaxis-mediated mechanisms and downregulated activation markers such as the expression of *Trem2*, which have previously been described to be regulated by the T_REG_-cytokine IL-10 (Shemer et al., 2020). These previous and our own results here clearly show the direct role of IL-10 in modulating microglial function. Likewise, using adult human microglial cells co-culture with T lymphocytes others demonstrated an enrichement of IL-10 secretion upon direct cell-cell contact (Chabot et al., 1999). In addition, we previously reported using whole genome sequencing that intracerebroventricular injection of IL-10 is sufficient to modulate the neuroinflammatory response after experimental stroke (Liesz et al., 2014).

An important caveat and potential key reason for the so far still pending success in harnessing the therapeutic function of IL-10 is its short half-life (less than 1 hour) and limited bioactivity after *in vivo* administration as a recombinant protein (Le et al., 1997; Saxena et al., 2015). Moreover, the systemic IL-10 application can have considerable and unforeseen side-effects due to the potentially divergent function of IL-10 on inflamed and homeostatic tissue, including direct effects on neurons, astrocytes, endothelial cells and other cellular constituents of physiological brain function (Saraiva et al., 2019). Therefore, we aimed to take a different approach for the localized and sustained production of IL-10 at the inflamed peri-lesional brain parenchyma. For this we took advantage of the potent capability of T cells to be specifically recruited and accumulated to the ischemic lesion site in order to deliver IL-10 from genetically engineered IL-10-overexpressing T cells (Heindl et al., 2021; Llovera et al., 2017). We demonstrated that IL-10 overexpression by this approach substantially modulated microglia function by down-regulation of microglial gene signature associated with phagocytosis of synapses correlating with functional recovery after stroke. Interestingly, eTc-IL10 cells did not exclusively invade the injured brain, but were also located in the meningeal compartment and could additionally contribute to functional recovery by resolving inflammation at these broder structures or providing IL-10 to the brain parenchyma along CSF flow. This concept is in accordance with previous observations of meningeal immune cell accumulation after stroke (Benakis et al., 2018) and that meningeal T cell-derived cytokines may enter the brain via CSF flow and paravascular spaces (Iliff et al., 2012).

An important finding in this study was the observation that IL-10 overexpression in T cells modulated microglial genes involved in the complement pathway, phagocytosis and synaptic pruning, and was associated with a better functional outcome after stroke. Complement factors are localized to developing CNS synapses during periods of active synapse elimination and are required for normal brain wiring (Schafer et al., 2012). Inactive synapses tagged with complement proteins such as C1q may be eliminated by microglial cells. Likewise in the mature brain, early synapse loss is a hallmark of several neurodegenerative diseases (Stephan et al., 2012). Indeed, complement proteins are profoundly upregulated in many CNS diseases prior to signs of neuron loss, suggesting mechanisms of complement-mediated synapse elimination regulated by microglia potentially driving disease progression (Stephan et al., 2012) and stroke recovery. It is therefore conceivable that T cells overexpressing IL-10 down-regulate the complement system in microglia and prevent excessive elimination of synapse and consequently protect against neuronal dysfunction. This is particularly of interest because microglia effector function have not only been associated with inflammatory neurodegenerative processes, but recently also been shown to be neuroprotective (Szalay et al., 2016) by tightly monitoring neuronal status through somatic junctions (Cserép et al., 2020). Microglia interact with the extra-neuronal space by not only regulating the elimination of existing synapses but also by modifying the extracellular matrix to enable efficient synaptic remodelling (Zaki and Cai, 2020). Accordingly, we found T cell-dependent regulation of several microglial genes that can mediate such extracellular matrix modifications involved in phagocytosis and proteases (*Clstn1* and *Mmp12*,cathepsins and MMPs, respectively).

Taken together, we have been able to demonstrate that brain-invading T cells can specifically “fine-tune” the transition of microglial to a reactive state. We postulate that the development of engineered T cells could have important translational implication by targeting a specific effector function of microglia with a relevant impact on the chronic progression of stroke pathobiology.

## Materials and Methods

### Animal experiments

All animal procedures were performed in accordance with the guidelines for the use of experimental animals and were approved by the respective governmental committees (Regierungspraesidium Oberbayern, the Rhineland Palatinate Landesuntersuchungsamt Koblenz). Wild-type C57BL6/J mice were purchased from Charles River, Rag-1^-/-^ mice (NOD.129S7(B6)-Rag-1tm1Mom/J) and eGFP-reporter mice (C57BL/6-Tg(CAG-EGFP)131Osb/LeySopJ) were bred and housed at the animal core facility of the Center for Stroke and Dementia Research (Munich, Germany). All mice were housed with free access to food and water at a 12 h dark-light cycle. Data were excluded from all mice that died during surgery. Animals were randomly assigned to treatment groups and all analyses were performed by investigators blinded to group allocation. All animal experiments were performed and reported in accordance with the ARRIVE guidelines (Kilkenny et al., 2011).

### Permanent distal middle cerebral artery occlusion model

Permanent coagulation of the middle cerebral artery (MCA) was performed as previously described (Llovera et al., 2014). Briefly, animals were anesthetized with volatile anesthesia (isofurane in 30%O_2_/70%N_2_O) and placed in lateral position. After a skin incision between eye and ear, the temporal muscle was removed and the MCA identifed. Then, a burr hole was drilled over the MCA and the dura mater was removed. The MCA was permanently occluded using bipolar electrocoagulation forceps. Permanent occlusion of the MCA was visually verifed before suturing the wound. During the surgery, body temperature was maintained using a feedback-controlled heating pad. Mice that developed a subarachnoid hemorrhage during surgery were excluded from the analysis.

### Cylinder test

To evaluate forepaw use and asymmetry, the cylinder test was performed two days prior to stroke (baseline) and day 2 and day 4 post stroke. Mice were placed in a transparent acrylic glass cylinder (diameter 8cm; height: 25 cm) in front of two mirrors and videotaped. To assess independent forelimb use, contact with one forelimb (left and right forelimbs) during full rearing and landing performance of mice were scored by frame-to-frame analysis of recorded videos. Mice with forepaw preference at baseline (absolute value difference between right and left forepaws > 10) were excluded from the analysis. All rearing movements during the trial were counted and used as indication of the animal’s overall activity.

### Intra cisterna magna injection

Mice were anesthetized with isofurane in 30%O_2_/70%N_2_O and fixed in a stereotaxic frame by the zygomatic arch, with the head slightly tilted to form an angle of 120° in relation to the body. A small incision was made at the nape of the neck between the ears to expose the neck muscles, which were bluntly dissected to expose the cisterna magna (CM). Cannulas composed of a glass capillary (ID, inner diameter 0.67 mm; OD, outside diameter, 1.20 mm) attached to a polyethylene tubing (ID 0.86 mm, OD 1.52mm, Fisher Scientific UK Ltd.) were used to perform the CM injections. Glass capillaries were sharpened using a flaming micropipette puller (P-1000, Sutter Instrument GmbH), filled with 10 μl of the cell suspension diluted in artificial CSF (aCSF: 126 mM NaCl, 2.5 mM KCl, 1.25 mM NaH_2_PO_4_, 2 mM Mg_2_SO_4_, 2 mM CaCl_2_, 10 mM glucose, 26 mM NaHCO_3_; pH 7.4 when gassed with 95% O_2_ and 5% CO_2_) and fixed to the micromanipulator arm of the stereotaxic. Cell suspension was injected into the CM at a rate of 2 μl/min. At the end of the injection mice are sutured and allowed to recover in a preheated awake cage for 1 h, after which they are returned to the animal husbandry.

### Infarct volume quantification

Mice were deeply anesthetized 5 days after stroke induction and transcardially perfused with 20 ml saline. Brains were removed, frozen immediately on powdered dry ice and stored at −20 °C until use. For infarct volumetry, brains were serially sectioned (400 μm intervals, 20 μm thick) and stained for cresyl violet (CV) as previously described (Llovera et al., 2014). CV stained sections were scanned at 600 dpi on a fatbed scanner (Canon). Direct infarct measurement was used after validating the absence of edema at the investigated time point. The total infarct volume was measured with ImageJ and determined by integrating measured areas and distances between sections.

### Immunohistochemistry and confocal microscopy

Microglia morphology analysis was performed on brain coronal sections as previously described (Heindl et al., 2018). Briefly, mice were perfused with 4% paraformaldehyde (PFA) and brains were post-fixed overnight and placed in sucrose for dehydration. Then, free floating 100 μm coronal sections were stained for microglia with 1:200 anti-Iba1 (rabbit, Wako, #019-19741). Nuclei were stained using 4’,6-Diamidin-2-phenylindol (DAPI, Invitrogen, #D1306) and images were acquired at a distance of 900 μm from the border of the lesion in layer 4 (ipsilateral) and the homotypic contralateral region using a Zeiss confocal microscope with 40x magnification (objective: EC Plan-Neofluar 40x/1.30 Oil DIC M27) with an image size of 1024 × 1024 pixel, a pixel scaling of 0.2 × 0.2 μm and a depth of 8 bit. Confocal-images were collected in Z-stacks with a slice-distance of 0.4 μm. Morphological features of microglia were acquired using a fully automated analysis as previously described (Heindl et al., 2018).

### Fluorescent In Situ Hybridization (FISH)

Single-molecule fluorescence in situ hybridization (FISH) was performed using the RNAscope Multiplex Fluorescent Reagent Kit v2 (Advanced Cell Diagnostics) by the manufacturer’s protocols. Briefly, free floating 100 μm coronal brain sections (Fig. 3e) or 20 μm cryo-sections (Fig. 4g) were first dried, washed, and then incubated in RNAscope hydrogen peroxide. Antigen retrieval and protease treatment were performed as per protocol. Sections were then incubated with the probe mix (C2-*Trem2* and C1-*Cx3cr1*) for 2 h at 40°C and then immediately washed with wash buffer. Next, sections were incubated with RNAscope Multiplex FL v2 AMP1, AMP2, and AMP3, and then probes were counterstained with TSA Plus Cy3 for C1-*Cx3cr1*, TSA Plus Cy5 for C2-*Trem2*. For microglia identification (Fig. 3e), slides were incubated in blocking at RT for 1 h before overnight incubation at 4°C with the primary rabbit anti-P2Y12 receptor antibody (1:200, AnaSpec #AS-55043A) and labelling for 1 hour with the secondary antibody AF488 goat anti-rabbit, (1:200, Invitrogen #A11034). Finally, sections were stained with DAPI (Invitrogen) and mounted with fluoromount medium (Sigma). smFISH-stained RNA molecules were counted only within the DAPI staining of the cell; a cell was considered *Cx3cr1*-positive when more than four *Cx3cr1* puncta were present.

### Whole skull immunofluorescence

Rag-1^-/-^ mice were anesthetized with isoflurane and perfused transcardially with ice-cold PBS followed by 4% PFA. After removing the mandibles, skin and muscles were carefully detached from the skull (http://www.nature.com/protocolexchange/protocols/3389). The skull decalcification was performed as previously described (Benakis et al., 2016). Coronal skull sections (20 μm) were stained with GFP-booster Atto647N (1:500, ChromoTek GmbH) to visualize eGFP-labelled T cells. Sections were counterstained with DAPI (Invitrogen) to visualize cell nuclei and observed by confocal laser microscopy (Leica SP5).

### In vitro T cell polarization

Single-cell suspensions were generated from spleen, inguinal, axial, brachial, and mandibular lymph nodes of C57BL/6 or b-actin-EGFP mice by passing the tissue through a 70-μm cell strainer. Naive CD4+ T cells were obtained by pre-enrichment using an “untouched” CD4+ T Cell Isolation Kit (Miltenyi Biotec) with subsequent flow cytometric analysis (CD4+ [clone RM4-5, 0.5 ng/μL], CD44low [clone IM7, 2 ng/μL], CD62Lhigh [MEL-14, 0.8 ng/μL]). Cells were seeded at a density of 300,000 or 400,000 cells/ well in a flat-bottom 96-well plate and stimulated with plate-bound anti-CD3 and anti-CD28 Abs (0.5 μg/mL or 2 μg/mL anti-CD3 [clone 145-2C11] for T_REG_ and T_H1_, respectively and 2 μg/mL anti-CD28 [clone 37.51]. Different mixtures of cytokines and mAbs were added to RPMI (supplemented with 10% FCS, 50 μM β-mercaptoethanol, 50 U/mL penicillin, 50 μg/mL streptomycin, 1% GlutaMAX™ and 1% N-2-hydroxyethylpiperazine-N-2-ethane sulfonic acid [Gibco® HEPES]) and used as follow: T_H1_ conditions with anti–IL-4 (10 μg/mL, BioXCell, #BE0045) and IL-12 (10 ng/mL, BioLegend, #577002); T_REG_ conditions: anti–IL-4 (10 μg/mL, BioXCell, #BE0045), anti-IFN-γ (10 μg/mL, BioXCell, #BE0055) and TGFβ (3ng/mL, BioLegend, #580702). After 2 days in culture, cells were split into 2 new 96-well plates and incubated with freshly prepared supplemented RPMI media with IL-2 (10 ng/ml, BioLegend, #575402). Cells were cultured for a total of 5 days before injection. Quality control was performed on day 4 to assess the percentage of T cell expressing Tbet (clone 4B10, 2 ng/μL) (T_H1_) or FoxP3 (clone FJK-16s, 2 ng/μL) (T_REG_) (Supplemental Fig. 3a). One million differentiated T cells were resuspended in sterile aCSF and injected into the CM in Rag1^-/-^ recipient mice 24 h after dMCAO induction.

### IL-10 overexpression in naïve T cells

Engineered T cells overexpressing IL-10 (eTc-IL10) were generated by transfection of naïve T cells with an IL-10 plasmid (pRP[Exp]-TagBFP2-CMV>mIl10[NM_010548.2]) designed and prepared by VectorBuilder (Supplementary Fig. 3c). First, splenocytes were isolated from C57BL/6 mice (male, 6-12 weeks old) and enriched using a CD4+ T Cell Isolation Kit (Miltenyi Biotec, No:130-104-453). Quality control was performed by flow cytometry (CD4+ [clone RM4-5, 1:25], CD44low [clone IM7, 1:25], CD62Lhigh [clone MEL-14, 1:25]). Cells were resuspended in RPMI (supplemented with 10% FBS, 50 μM β-mercaptoethanol, 50 U/mL penicillin, 50 μg/mL streptomycin, 1% GlutaMAX, 1% HEPES and 10 ng/mL IL-2). To induce CD4+ cells to enter the cell cycle for efficient DNA uptake, 4×10^5^ cells/well were seeded in a flat-bottom 96-well plate containing bound anti-CD3 (2 μg/mL, clone 145-2C11) and anti-CD28 (2 μg/mL, clone 37.51) for 48 h. After 48 h stimulation, 1.5×10^6^ cells multiplied by the number of mice to be injected were transfected with the pIL-10 vector (1×10^6^ cells/1.5 μg pIL-10 DNA per cuvette) using the Mouse T Cell Nucleofector Kit (Lonza, No: VPA-1006) with Nucleofector II Device (program X-100). Once electroporated, cells were diluted with conditioned RPMI from the 48 h stimulation and fresh supplemented RPMI (1:1) and seeded in a 12-well plate (1 cuvette of cells/ well). 24 h post transfection, cells and supernatant were collected. Supernatant was used to confirm IL-10 secretion by ELISA (Supplementary Fig. 3c) and cells were collected for intra CM injection (1×10^6^/mouse).

### ELISA

Secreted IL-10 was determined by ELISA as per the manufacturer’s protocol (Mouse IL-10, Invitrogen, No: 88-7105-88). The color reaction was measured as OD450 units on a Bio-Rad iMark microplate reader. The concentration of supernatant IL-10 was determined using the manufacturer’s standard curve over the range of 32-4’000 pg/mL.

### Flow cytometry

For differentiation of live and dead cells we stained cells with the Zombie Violet Fixable Viability Kit according to the manufacturer’s instructions (BioLegend). For surface marker analysis, cell suspensions were adjusted to a density of 0.5×10^6^ cells in 50 μl FACS buffer (2% FBS, 0.05% NaN3 in PBS). Nonspecific binding was blocked by incubation for 10 min at 4 °C with anti-CD16/CD32 antibody (Biolegend, clone 93, 5 ng/ μL) antibody and stained with the appropriate antibodies for 15 min at 4 °C. The following antibodies were used for extracellular staining: CD45 (clone 30F-11, 0.5 ng/μL), CD4 (clone RM4-5, 0.5 ng/μL), CD11b (clone M1/70, 0.6 ng/μL), CD19 (eBio1D3, 0.6 ng/μL), B220 (clone RA3-6B2, 0.32 ng/μL), CD3ε (clone 145-2C11, 2 ng/μL), CD8a (clone 53-6.7, 2 ng/μL), CD62L (clone MEL-14, 0.8ng/μL) from Thermofisher. For intracellular cytokine staining, cells were restimulated for 4 hours with PMA (50ng/ml, Sigma), ionomycin (1 μM, Sigma) and brefeldin A (1 μl for ~10^6^ cells/mL). Cells were then stained for surface markers as detailed below, fixed and permeabilized using Fixation and Permeabilization Buffers from eBiosciences following the manufacturer’s instructions. Briefly, cells were fixed for 30 min at 4 °C (or RT for FoxP3), washed with permeabilization buffer and incubated for 30 min with the appropriate antibodies in permeabilization buffer at 4 °C (or RT for FoxP3). The cells were stained with the transcription factors FoxP3 (clone FJK-16s, 2 ng/μL) and T-bet (clone 4B10, 2 ng/μL) or IFN-γ (clone 4S.B3, 2 ng/μL). Cells were washed with FACS buffer, resuspended in 200 μl of FACS buffer and acquired using a BD FACSverse flow cytometer (BD Biosciences, Germany) and analyzed using FlowJo software (Treestar, US). Isotype controls were used to establish compensation and gating parameters.

### Nanostring analysis

The ipsilateral hemispheres were lysed in Qiazol Lysis Reagent and total RNA was extracted using the MaXtract High Density kit with further purification using the RNeasy Mini Kit (all Qiagen). 70 ng of total RNA per sample was then hybridized with reporter and capture probes for nCounter Gene Expression code sets (Mouse Neuroinflammation codeset) according to the manufacturer’s instructions (NanoString Technologies). Samples (6/condition) were injected into NanoString cartridge and measurement run was performed according to nCounter SPRINT protocol. Background (negative control) was quantified by code set intrinsic molecular color-coded barcodes lacking the RNA linkage. As positive control code set intrinsic control RNAs were used at increasing concentrations. Genes below the maximal values of the negative controls were excluded from the analysis. All gene counts were normalized (by median) and scaled (mean-centered and divided by standard deviation of each variable). Heatmaps were performed using the MetaboAnalystR package on normalized expression values. The regulated genes in microglia treated with T_H1_ or T_REG_ in comparison to vehicle treated microglia (CT) are represented in the volcano plots, genes with a P < 0.05 were color-coded. Significantly up-regulated genes with a FC > 3 which were specific to T_H1_ or T_REG_ (Venn diagram) were further used for pathway analysis using Cytoscape ClueGO (T_H1_/CT: 21 out of 26 genes, T_REG_/CT: 5 out of 10 genes (Bindea et al., 2009)).

### Microglia cell isolation for RNA sequencing

Mice were perfused transcardially with ice-cold saline containing Heparin (2U/mL). Brains were placed in HBSS (w/ divalent cations Ca^2+^ and Mg^2+^) supplemented with actinomycin D (1:1000, 1 mg/mL, Sigma, #A1410), and microglia was isolated with the Papain-based Neural Tissue Dissociation Kit (P) (# 130-092-628, Miltenyi Biotec B.V. & Co. KG) according to the manufacturer’s instructions. Cell suspension was enriched using 30% isotonic Percoll gradient. 1×10^3^-1.5×10^3^ live microglia cells from 3 mice per condition were sorted according to their surface marker CD45+CD11b+7-AAD negative (SH800S Cell Sorter, Sony Biotechnology) and proceed for 10x Genomics according to the manufacturer’s instructions (ChromiumTM Single Cell 3’ Reagent kits v2). For the Smart-seq2 platform, pools of 50 cells were sorted into 96 well plates, already filled with 4 μL lysis buffer containing 0.05% Triton X-100 (Sigma) and, ERCC (External RNA Controls Consortium) RNA spike-in Mix (Ambion, Life Technologies) (1:24000000 dilution), 2.5 μM oligo-dT, 2.5 mM dNTP and 2 U/μL of recombinant RNase inhibitor (Clontech) then spun down and frozen at –80 °C.

### Single cell data analysis

The CellRanger software (v2.0.0, 10X Genomics) was used for demultiplexing of binary base call (BCL) files, read alignment, and filtering and counting of barcodes and unique molecular identifiers (UMI). Reads were mapped to the mouse genome assembly reference from Ensembl (mm10/GRCm38).

Downstream data analyses were performed using the Scanpy API (scanpy v>=1.4 with python3 v>=3.5; (Wolf et al., 2018)). Details on analyses, selected thresholds and package versions are provided in available source scripts (See Code and Data availability). Outlier and low-quality cells were filtered if the fraction of mitochondria-encoded counts was greater than 10% or the total number of counts was greater than 48’000. Thresholds were selected upon visual inspection of distributions as recommended (Luecken and Theis, 2019). Genes expressed in less than 10 cells were excluded. Further, doublet cells as identified by the Scrublet algorithm (v0.2.1; (Wolock et al., 2019)) were excluded. Doublet scores and thresholds were determined for each sample separately. Raw counts of a cell were normalized by total counts neglecting highly expressed genes which constitute more than 5% of total counts in that cell. Then, counts were log-transformed (log(count+1)). These processed and normalized count matrices were used as input for all further analyses.

For the full data set and the microglia subset first a single-cell nearest-neighbor graph was computed on the first 50 independent principal components. Principle components were calculated using the 3000 most variable genes of the full data set as input. The UMAP algorithm (Becht et al., 2019) as used to obtain a two-dimensional embedding for visualization. Iterative clustering was performed with the Louvain algorithm (Blondel et al., 2008) as implemented in louvain-igraph (v0.6.1 https://github.com/vtraag/louvain-igraph) with a varying resolution parameter. Clusters were annotated using previously described marker genes and merged if expressing the same set of marker genes.

Trajectories from homeostatic to reactive microglia were inferred with partition-based graph abstraction (PAGA) (Wolf et al., 2018) and diffusion pseudotime (DPT) (Haghverdi et al., 2016) algorithms. First, clusters were grouped into two paths connecting the root and end cell cluster based on the computed cluster connectivities (PAGA), then, cells were ordered along these paths based on the random-walk based cell-to-cell distance (DPT). To capture processes specific to the path 2 trajectory in stroke-associated microglia, data was first subset to cells of path 2 and end cells clusters of stroke samples and gene expressed in less than 20 cells of the subset excluded. Then, gene sets were computed by clustering the 500 most varying genes using their pairwise-pearson correlation values as input and Ward’s hierarchical clustering method with euclidean distance (scipy python package v.1.5.4 (Virtanen et al., 2020)). One gene set with average correlation < 0.05 was excluded. Finally, to obtain an activation score per cell for a given gene set, cell scores were computed as described by (Satija et al., 2015) and implemented in Scanpy in the *tl.score_genes* functionality. Differential activation of gene sets between WT and Rag1^-/-^ samples was determined by a Wilcoxon rank-sum test. To identify genes differentially regulated along the inferred cellular trajectory a differential gene expression test (Welch t-test with overestimated variance) between the root and end cell cluster was performed for WT and Rag1^-/-^ samples separately. Non-overlapping, significantly changing genes (p-value <0.05 corrected for multiple testing with the Benjamin-Hochberg method) were considered as regulated specifically in WT and Rag1^-/-^ samples, respectively. Pathway enrichment of gene sets and differentially regulated genes was performed with the gseapy package (https://github.com/zqfang/GSEApy/) functionality of EnrichR (Xie et al., 2021).

### 10x Genomics data and code availability

Jupyter notebooks with custom python scripts for scRNA-seq analysis will be made available in a github repository upon publication (https://github.com/theislab/). 10X Genomics and Smart-seq2 data have been submitted to GEO and will be made available upon publication.

### Library preparation for Smart-seq2 platform

The 96-well plates containing the sorted pools were first thawed and then incubated for 3 min at 72°C and thereafter immediately placed on ice. To perform reverse transcription (RT), we added each well a mix of 0.59 μL H2O, 0.5 μL SMARTScribe™ Reverse Transcriptase (Clontech), 2 μL 5x First Strand buffer, 0.25 μL Recombinant RNase Inhibitor (Clontech), 2 μL Betaine (5 M Sigma), 0.5 μL DTT (100 mM), 0.06 μL MgCl2 (1 M Sigma), 0.1 μL Template-switching oligos (TSO) (100 μM AAGCAGTGGTATCAACGCAGAGTACrGrG+G). Next, RT reaction mixes were incubated at 42°C for 90 min followed by 70°C for 5 min and 10 cycles of 50°C 2 min, 42°C 2 min; finally ending with 70°C for 5 min for enzyme inactivation. Pre-amplification of cDNA was performed by adding 12.5 μL KAPA HiFi Hotstart 2x (KAPA Biosystems), 2.138 μL H2O, 0.25 μL ISPCR primers (10 μM, 5’ AAGCAGTGGTATCAACGCAGAGT-3), 0.1125 μL Lambda Exonuclease under the following conditions: 37°C for 30 min, 95°C for 3 min, 20 cycles of (98°C for 20 sec, 67°C for 15 sec, 72°C for 4 min), and a final extension at 72°C for 5 min. Libraries were then cleaned using AMPure bead (Beckman-Coulter) cleanup at a 0.7:1 ratio of beads to PCR product. Library was assessed by Bio-analyzer (Agilent 2100), using the High Sensitivity DNA analysis kit, and also fluorometrically using Qubit’s DNA HS assay kits and a Qubit 4.0 Fluorometer (Invitrogen, LifeTechnologies) to measure the concentrations. Samples were normalized to 160 pg/μL. Sequencing libraries were constructed by using an in-house produced Tn5 transposase (Picelli et al., 2014). Libraries were barcoded with the Illumina Nextera XT (FC-131-1096, Illumina) and pooled, then underwent three rounds of AMPure bead (Beckman-Coulter) cleanup at a 0.8:1 ratio of beads to library. Libraries were sequenced at 2×100 base pairs paired-end on Illumina HiSeq4000.

### Processing and analyses of Smart-seq2 data

BCL files were demultiplexed with the bcl2fastq software from Illumina. After quality-control with FastQC, reads were aligned using rnaSTAR (Dobin et al., 2013) to the GRCm38 (mm10) genome with ERCC synthetic RNA added. Read counts were collected using the parameter “quantMode GeneCounts” of rnaSTAR and using the unstranded values. From that point, Seurat R v.3 package was used (Stuart et al., 2019). Low-quality samples were filtered out from the dataset based on a threshold for the number of genes detected (min 5000, max 12500 unique genes/pool), percentage of mitochondrial genes (maximum: 1.5 %), percentage of ERCCs (maximum: 0.5 %) and number of reads (between 100k to 3M). 291 pools passed the quality-control. Gene counts were log normalized using the NormalizeData function of Seurat with a scale factor of 50,000. Dataset were scaled using the ScaleData function. The top 1000 most variable genes were considered for the rest of the analyses. The first 13 principal components were kept from the PCA. Differential expression analysis was performed using DESeq2 (Love et al., 2014) using the FindAllMarkers function of Seurat. Gene ontology analyses were performed using Metascape (Zhou et al., 2019) using the genes significantly differentially expressed at adjusted p-value of 5 %. Sample clustering was performed using the 1000 most variable genes. First, Pearson correlation coefficient was calculated between each sample. Then, hierarchical clustering was performed (hclust function from stats package) on the Euclidian distance between samples via the dist function from the stats package and using the ward.D2 method. Conversely, gene clustering was performed similarly starting from the transposed expression matrix. Gene ontology was then performed on each gene cluster using Metascape (Zhou et al., 2019). Figures were generated with ggplot2 package (Wickham, 2016).

### Statistical analysis

Data are expressed as mean ± s.e.m. and were analyzed by unpaired Student’s *t*-test (two-tailed) or one- or two-way ANOVA and post-hoc tests as indicated in the figure legends. Exclusion criteria are described in the individual method sections. The data for microglia morphology is shown as median ± interquartile range and statistical significance was tested using the Wilcoxon rank sum test with continuity correction and Bonferroni post-hoc correction for multiple testing in R (version 4.0.3).

## Author contributions

C.B.: conceptualization, formal analysis, investigation, methodology, supervision, visualization, writing – original draft, funding acquisition. A.S: investigation, methodology (stroke surgery, CM injection, T cell polarization), validation, supervision; S.H: data curation, formal analysis, investigation (microglia morphology and analysis); K.P: investigation, methodology (engineered T cells, ELISA), formal analysis; G.L: investigation, methodology (FISH staining and quantification), formal analysis and A.K: investigation, methodology (T cell polarization). S.T. and F.T. performed the 10x Genomics analysis. S.B-G and O.G. ran the Smart-seq2 platform and performed the bioinformatics analysis. A.K. and A.P. performed the *in vitro* polarization of regulatory T cells. A.L.: funding acquisition, supervision, project administration, conceptualization, manuscript review, and editing.

## Acknowledgments

The authors thank Kerstin Thuβ-Silczak and Dr. Monica Weiler for technical support and Michael Heide and Oliver Weigert for support at the DKTK Nanostring core facility. Graphical schemes in Figures 1, 4 and Supplementary Figure 3 were created with BioRender.com. The study was supported by the European Research Council (ERC-StG 802305) and the German Research Foundation (DFG) under Germany’s Excellence Strategy (EXC 2145 SyNergy – ID 390857198), through SFB TRR 274 and under the DFG projects 405358801 (to A.L.), 418128679 (to C.B.) and PE-2681/1-1 (to A.P.). The authors declare no competing financial interests.

---

**Supplementary Fig. 1.**
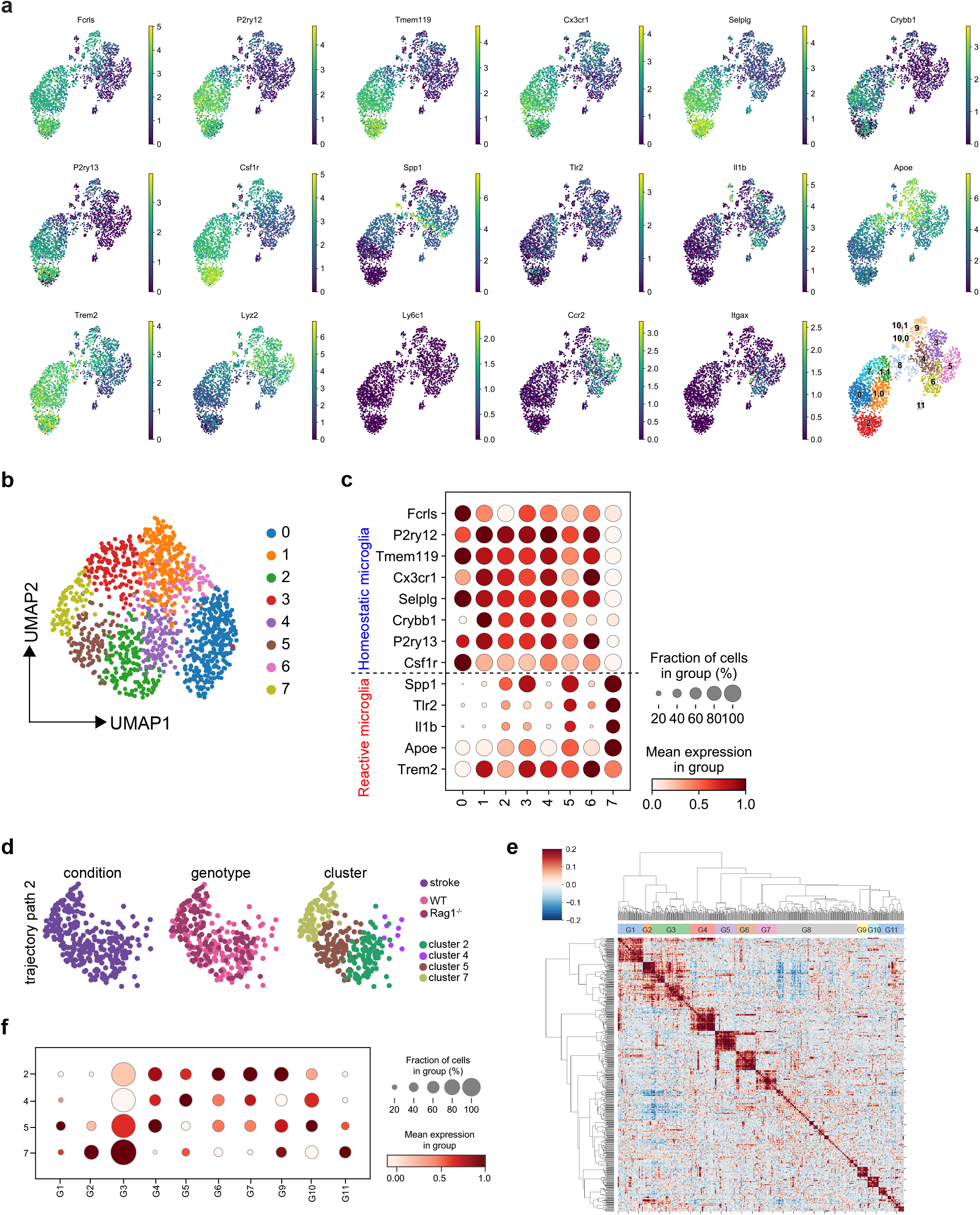
Transcriptomic analysis of microglia isolated from WT and Rag1^-/-^ in naïve and stroke conditions. **a** UMAP plots showing expression of known CD45+CD11b+ myeloid cell marker genes and Louvain-clusters. **b,c** Manifold and clustering of microglia: **b** UMAP plots indicate Louvain-clusters of microglial cells. **c** Dot plots show gene expression distribution of marker genes split by Louvain-cluster, and their grouping into homeostatic (top) and reactive (bottom) microglia. **d–e** Microglia gene set expression in stroke condition. **d** UMAP plots in path 2 and end cell clusters colored-coded by stroke condition (left), genotype (middle) and clusters (right). **e** Gene-gene correlation map of the correlated and anti-correlated genes extracted from clusters 2, 4, 5 and 7 identified 11 gene clusters. Due to low correlation, G8 was excluded of further analysis. **f** Dot plots show gene set expression distribution (G1 to G11) of the stroke cell clusters (2, 4, 5, 7).

**Supplementary Fig. 2.**
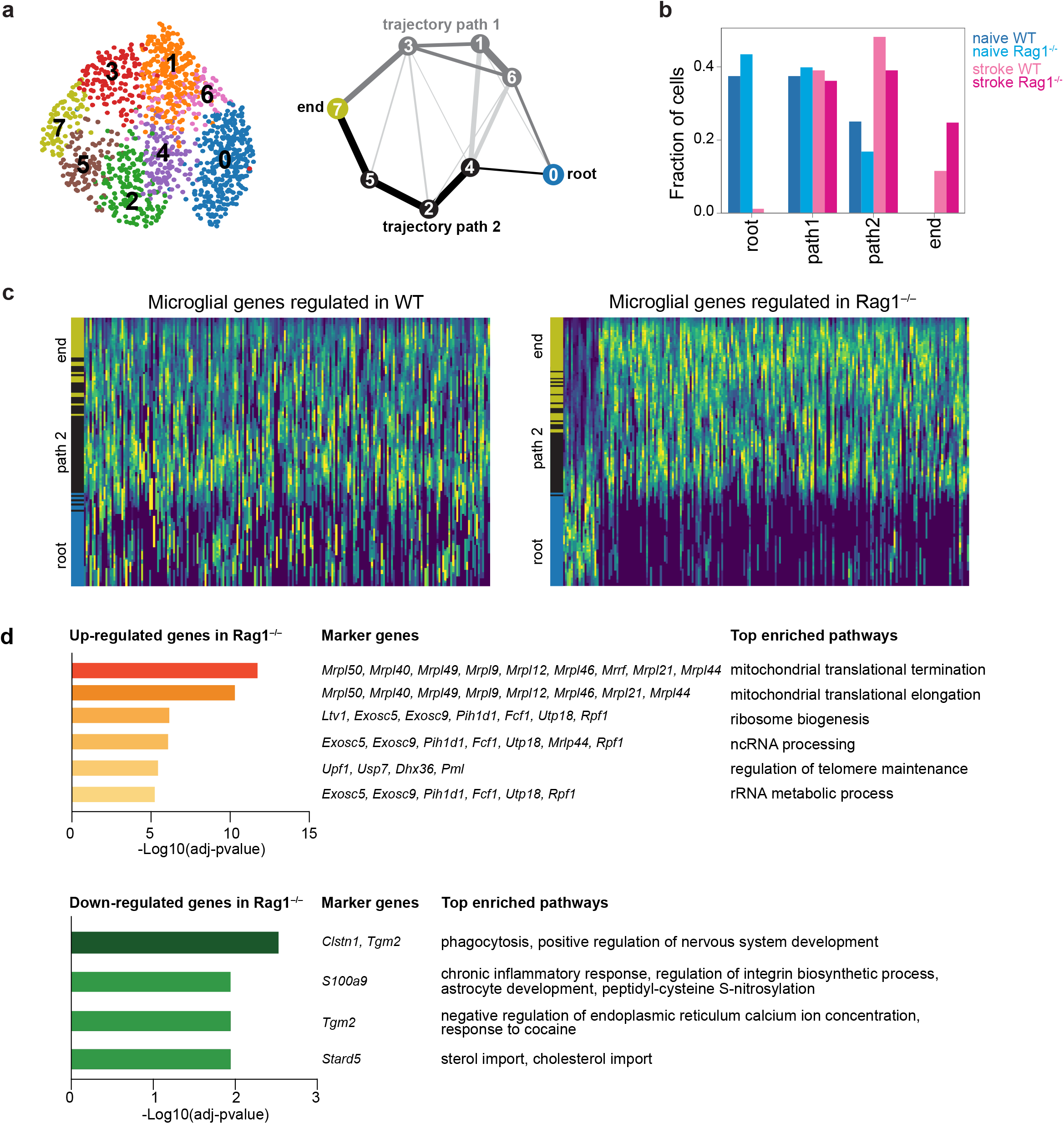
Microglia single cell trajectory inference in WT and Rag1^-/-^ naïve and stroke conditions. **a,b** Microglia single cell trajectory inference describes the evolution of microglia activation from naïve to stroke condition. **a** Partition-based graph abstraction graph (PAGA, right) shows cluster connectivity of Louvain-clusters (left) with a threshold of 0.1. Nodes represent subsets, and thicker edges indicate stronger connectedness between subsets. The trajectories path 1 and 2 are selected based on the connectivity (edge width) in the PAGA graph after the root and end paths were defined based on marker gene expression. Clusters along the paths from root-to-end were merged since these are consecutive. Trajectory path 1 (dark grey): root (cluster 0), path 1 (cluster 6, 1, 3 merged), end (cluster 7); trajectory path 2 (black): root (cluster 0), path 2 (cluster 4, 2, 5 merged), end (cluster 7). **b** Barplot shows cell frequencies split by condition in trajectories path 1 and path 2. **c,d** Differential microglial gene expression analysis between root and end clusters along trajectory path 2. **c** Heatmaps show scaled expression along path 2 trajectory of microglial genes specifically regulated in Rag1^-/-^ and compared in WT (left) and Rag1^-/-^ (right) in path 2 (Wilcoxon test, adjusted P < 0.05). For balanced representation of cells along the trajectory root and path 2, cells were randomly subset to 100 cells before plotting. **d** Top enriched pathways of microglial genes up- and down-regulated in Rag1^-/-^ mice.

**Supplementary Fig. 3.**
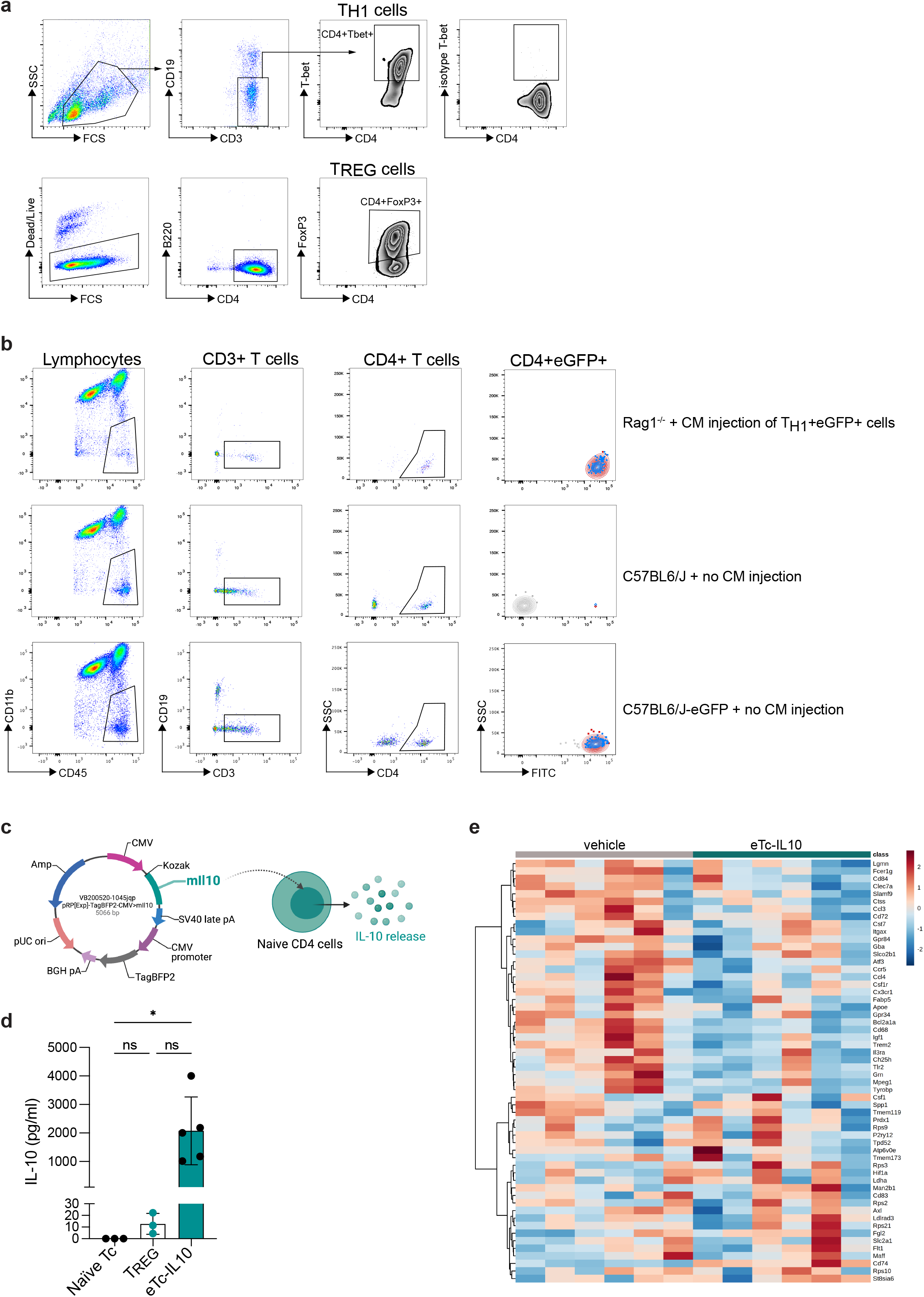
T cell polarization in vitro and IL-10 plasmid construct. **a** CD4 naïve T cells were polarized *in vitro* towards T_H1_ (top row, CD4+Tbet+) or T_REG_ (bottom row, CD4+FoxP3+). **b** Gating strategy of CD4^+^eGFP^+^ cells analyzed by flow cytometry in the ipsilateral (red) and contralateral (blue) hemispheres (CD45^+^CD11b^-^CD19^-^CD3^+^CD4^+^FITC^+^) in Rag1^-/-^ (top row), C57BL6/J (no cell injection, mid-row) and in C57BL6/J actin-eGFP mice (no cell injection, bottom row). **c** Plasmid expression vector pIL-10 was constructed by inserting 537bp IL-10 cDNA under a CMV promoter with a BFP2 tag and ampicillin resistance. **d** IL-10 concentration measured by ELISA in naïve T cells, T_REG_ cells and naïve T cell transfected with pIL-10 (eTc-IL10); ns, non significant; *, P < 0.05. **e** Heatmap representation of brain gene expression between vehicle and eTc-IL10 treated mice selected from the known disease-associated microglial genes (Keren-Shaul et al., 2017).

